# Assessing the Nucleotide-Level Impact of Spaceflight Stress using RNA-Sequencing Data

**DOI:** 10.1101/2022.12.01.518235

**Authors:** Montana S. Knight, Colleen J. Doherty, Dahlia M. Nielsen

## Abstract

Understanding the effects of space radiation and microgravity on DNA integrity is critical to assess the impact of long-term spaceflight. However, studying spaceflight’s effect on terrestrial life is difficult. NASA created GeneLab, a public Omics database for spaceflight-related data, to help combat these limitations. While GeneLab has very few DNA-based data sets, transcriptome information is abundant. This study used RNA-Seq data from GeneLab to examine DNA sequence variants linked to spaceflight stress exposure. More mutations were observed in spaceflight samples than in the ground control samples. This increase in variants was not reduced in samples grown under artificial gravity in space, suggesting that microgravity did not significantly affect the amount of DNA damage in this experiment. There was also an increase in transversion mutations, consistent with known forms of radiation-induced damage. This work demonstrates that RNA-Seq data is a useful resource for evaluating DNA damage from spaceflight and provides a baseline for the types of mutations that could be detected.

## Introduction

It is important to understand how spaceflight impacts terrestrial life to ensure astronauts’ health and safety and optimize future space travel^1-3^. Two major environmental stressors during spaceflight, microgravity and space radiation, are known to impact terrestrial life. Microgravity has been linked to muscle atrophy in astronauts, disruptions to DNA repair enzymes, and impacts on cellular proliferation^4-7^. Space radiation comes in two forms: galactic cosmic rays (GCR) and solar radiation^8-10^. GCRs consist of high-energy protons and high-energy charge (HZE) nuclei^8,9^. Solar radiation is electromagnetic radiation from the sun. It is sporadic but powerful. Solar radiation follows a cycle, making it variable over days and even years and thus more difficult to study^8^.

The best way to understand space radiation’s impact on terrestrial life is through experiments in true spaceflight. Such space-based experiments have demonstrated that heavy ion radiation impacts DNA. For example, chromosomal aberrations were observed in HeLa cells exposed for 40 days on the Russian MIR space station^4^. Damage was also observed in cells exposed for as little as nine days on the Space Shuttle^4^. Long-term space radiation exposure increases astronauts’ risk of cancer, cataracts, and neurological disorders^11,12^. In plants, spaceflight has demonstrated links to changed alternative splicing patterns, epigenetic patterns, and germination rates^7,13-15^. However, these studies fail to separate microgravity and non-microgravity-related spaceflight stress to properly investigate the potential impact of space radiation. These results indicate that more research is needed to understand the full scope of space radiation and capture its variable nature to assess its associated damages to terrestrial life better.

Experiments performed in space to observe the effects of true spaceflight on terrestrial life are informative but expensive and difficult to perform due to limited resources^16,17^. One alternative to space-based studies is to simulate the spaceflight conditions on Earth^18-20^. Microgravity can be simulated with drop towers, 3D clinostats, and random positioning machines^19^. Space radiation can be evaluated at facilities like the NASA Space Radiation Laboratory in Upton, NY^18^. Damage from radiation can also be evaluated with high-altitude balloon flights. These simulation procedures are good ways to approximate the impact of spaceflight^4,20^. One study observed DNA structural variants in *Arabidopsis thaliana* seeds exposed to space radiation on a high-altitude balloon over Antarctica, demonstrating such balloons as a practical way to study radiation without entering low earth orbit (LEO)^21^. Other types of radiation, such as ultraviolet, are easily studied on Earth. Previous research showed solar radiation experienced on Earth (which contains multiple types of UV radiation) is linked to increased transversion mutations^22^. UVA radiation was demonstrated to lead to a significant increase in transversion mutations in mice^23^. Other studies have pointed to an increase in radical oxygen species due to solar radiation. Further, DNA damage from these radicals was associated with a rise in G/C transversion mutations^22^. These approximations for both space radiation and microgravity conditions are useful. However, it is important to validate their findings in spaceflight to confirm if the specific radiation experienced during spaceflight has similar impacts on terrestrial life.

The prohibitive nature of ISS experiments makes it valuable to reevaluate data from previous spaceflight experiments to gain additional insights^24^. NASA created a public Omics data repository, GeneLab, to facilitate the reevaluation of ISS-collected data. GeneLab contains many Omics datasets from true spaceflight experiments. One drawback is that GeneLab is heavily biased toward transcriptome datasets, most of which were intended for RNA analysis. However, data from RNA-Seq experiments can be further analyzed to gain insights about DNA damage without having to do an additional spaceflight experiment.

This study uses GeneLab data to identify novel impacts of spaceflight stress. We demonstrate that identifying and examining variants in RNA-seq data from *Arabidopsis thaliana* samples grown aboard the ISS provides a proxy for estimating DNA damage from spaceflight. The number of single nucleotide variants in ISS-grown *A. thaliana* samples was identified and compared to the ground control samples. As expected, spaceflight samples resulted in a higher number of variants. Additionally, variant composition in each sample was examined, and spaceflight was associated with a higher number of transversion mutations. This aligns with previous studies examining the impact of radiation on the nucleotide level^22,23^. Thus, this study demonstrates RNA-Seq data can be utilized to evaluate the effects on DNA.

## Materials and Methods

### Datasets and Variant Calling Preparation

GeneLab’s data was examined for species and assay type composition on August 29, 2022. The data was downloaded from NASA’s GeneLab (GLDS-223)^25^. Raw fastq files were analyzed for any issues with fastqc^26^. Fastx_trimmer was used to trim the first thirteen and last five nucleotides of each read in each sequence file^27^. Fastq_quality_filter was then used to filter out base calls with a phred-quality score lower than 30^27^. Fastqc was again used to examine the quality of the newly filtered fastq files before aligning the reads to the Arabidopsis genome. Trimmed and filtered fastq reads were mapped to the TAIR10 genome using STAR in 2-pass mode^28-30^. Alignment files were evaluated with Picard’s “CollectAlignmentSummaryMetrics”. This served as a quality control check of the alignments^31^. STAR’s alignment files were then prepared for variant calling according to the somatic variant calling pipeline described in Knight et al. 2022^32-33^. The pipeline is available on GitHub (https://github.com/montana-knight/Calling-Induced-Variants-with-RNA-Seq-Data).

### Generating a Panel of Normals

Mutations that likely existed in the parental genotypes of both the spaceflight and ground control samples but differed from the reference sequence needed to be identified, so they were not included in downstream analysis. Later, this list of likely inherited mutations, called the Ultimate Panel of Normals (PON) is used as the “known list of population variants” for BaseRecalibrator and the “PON” for Mutect2^35^. To find these potential inherited mutations, four sets of likely inherited mutations in the Arabidopsis samples were made, one for each combination of tissue type and genotype (WT + root, WT + shoot, transgenic + root, transgenic + shoot). This was to ensure inherited mutations different between tissue or genotype type were accounted for. The sets of variants were created according to the pipeline outlined in Knight et al. 2022^32-33^. HaplotypeCaller^34^ was run with a heterozygosity value of 0.0001 to accommodate for the highly inbred nature of Arabidopsis. Mutect2^35^ was run in “tumor-only” mode with a max number of haplotypes of 2 (also to accommodate the low level of heterozygosity in Arabidopsis). The two sets of joint variant calls from HaplotypeCaller^34^ and Mutect2^35^ were combined, so variants called by either tool and present in at least three samples were considered likely inherited variants. This final list of likely inherited mutations was the Ultimate PON and was used as the “known list of population variants” for BaseRecalibrator and the “PON” for Mutect2^35^.

### Variant Calling and Filtering

Variants were called using Mutect2 in “tumor-only” mode with the same parameters used to create the PON. Variants were first filtered using GATK’s VariantFiltration^35^. Variants listed in the Ultimate PON were filtered out to remove likely inherited mutations. Variants with read depths lower than 10 and log odds lower than 6 were filtered out. Variants that clustered in groups of 3 or more within a span of 35 nucleotides were also filtered out. Next, variants were filtered based on the deduplication protocol in the Knight et al. 2022 pipeline^32-33^. Variants were also removed if they were in regions with increased opportunity for sequencing error, in lowly expressed genes (less than 10 pseudo-counts), or not expressed in all the samples. Regions with increased risk of sequencing errors were defined as a sequence of length three or more composed of the same nucleotide. Variants called in these sequences (ex. AA”A”) were removed. This is a conservative approach to identifying high-confidence variants. Gene expression levels were determined by htseq-count using the alignment file from STAR 2-pass and the TAIR10 annotation file^28,36^. The final output from the variant calling and filtering steps is a variant composition file for each sample.

### Variant Calling Composition

A custom script was created to analyze each sample’s variant call file (VCF) and obtain information on the composition of the mutations. The script outputs the number of:

- Single nucleotide polymorphisms (SNPs)
- Indels (both insertions and deletions)
- Times the alternative SNP was an A, C, G, or T
- Transversion mutations
- Transition mutations
- Times the SNP was an A to C, G, or T variant
- Times the SNP was a C to A, G, or T variant
- Times the SNP was a G to A, C, or T variant
- Times the SNP was a T to A, C, or G variant

### Statistical Analysis

Final variant counts were analyzed in R with a generalized linear model. Spaceflight conditions, tissue type, and Arabidopsis genotype were all tested for their contribution to the results. Least square estimates were obtained for all treatment effects including the main effects of potential interactions. The variant composition was also examined using a generalized linear model with a binomial distribution. Proportions for each type of variant were examined, and treatment effects were tested for their influence on the results.

## Results

### GeneLab has a high proportion of gene expression data

The purpose of this study was to examine the impact of spaceflight stress at the nucleotide level using publicly available data on GeneLab, a data repository set up by NASA for data from spaceflight-related experiments. It currently holds data from 385 studies (August 29, 2022). The data encompass actual and simulated spaceflight conditions, with 54.81% of the datasets having a true spaceflight condition component (Fig. 1 and Supplementary Fig. S1). We examined the composition of the different data types to assess the kinds of analysis necessary to achieve our objective.

**Figure 1.**
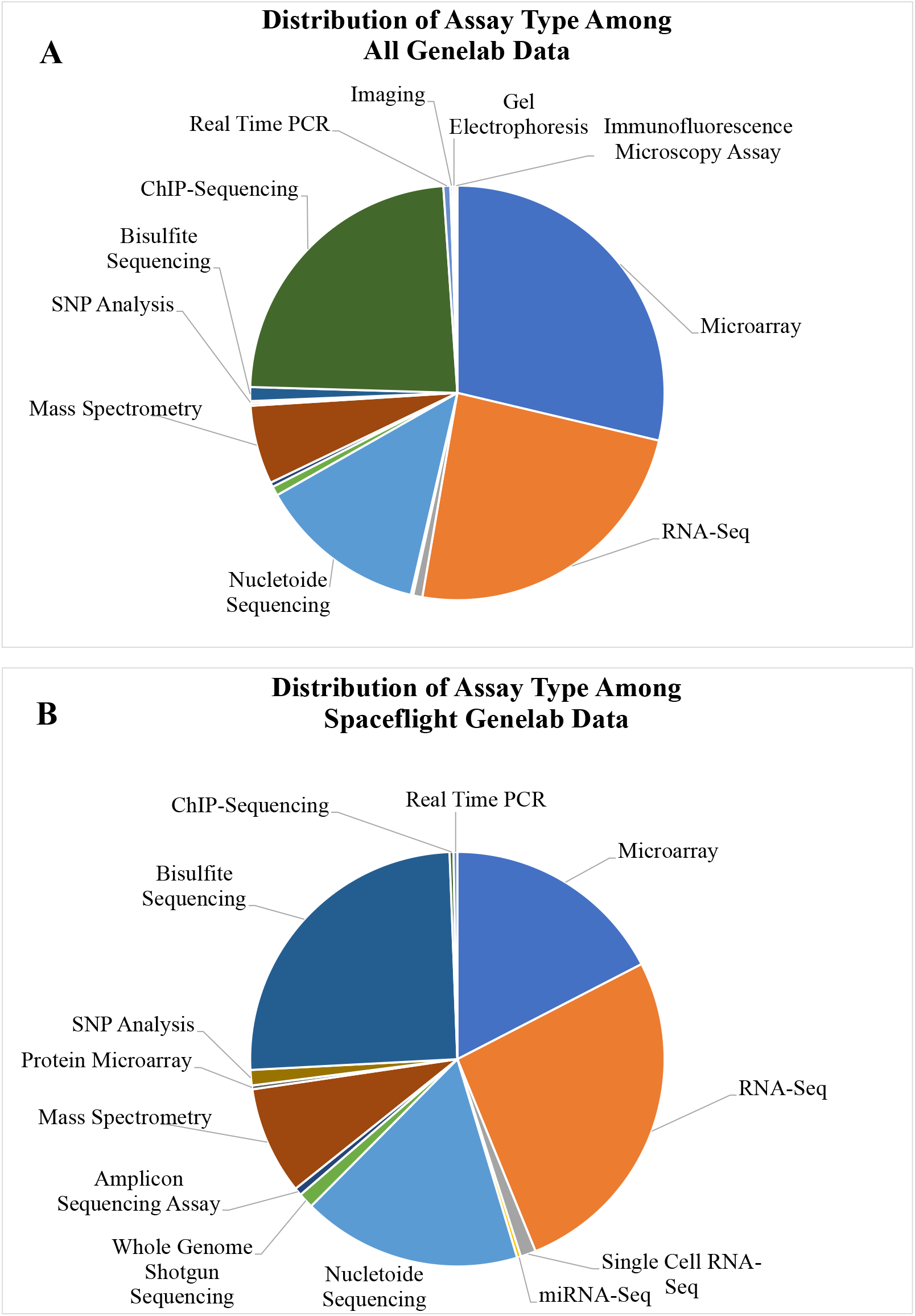
Types of experiments with data available on GeneLab. (Data Accessed August 29, 2022). GeneLab experiments encompass many different biological assays. (A) The distributions of these experiments are slightly different between all GeneLab experiments and (B) those experiments with a spaceflight component. Transcriptomic information obtained through microarray and RNA-Seq are the most common type of data on GeneLab.

GeneLab data includes many species. *Mus musculus* makes up 32.21% of all GeneLab datasets and 36.02% of all spaceflight datasets (Supplementary Fig. S1). Plants are another large section of the database, with 13.51% of all available datasets and 12.80% of spaceflight-specific datasets examining plants (Supplementary Fig. S1). Of this plant data, 92.31% of the data in all GeneLab sets and 88.89% of data in spaceflight-only sets are from Arabidopsis.

The composition of GeneLab’s assay types was also examined. 77.14% of studies have a transcription profiling component, demonstrating a major bias toward gene expression experiments. RNA-Seq data makes up 34.55% of all GeneLab data and 41.71% of the spaceflight data (Fig. 1). There are a total of 133 datasets from RNA-Seq experiments with a variety of species, including 76 sets from *M. musculus*, 23 sets from *Arabidopsis*, and seven from humans. Eighty-eight of these datasets are spaceflight-specific RNA-Seq datasets; 13 are from *Arabidopsis*, 53 are from *M. musculus*, and five are from humans. Less than a third of the spaceflight studies have a DNA component, and over half of the spaceflight experiments with nucleotide sequencing information are from single-cell organisms. No public spaceflight dataset on GeneLab from humans or plants with DNA sequence information is available to assess DNA damage. However, the abundance of RNA-Sequencing data provides an opportunity to analyze transcripts for variants and gain insight into how spaceflight impacts terrestrial life at the nucleotide level. Therefore, we examined the potential to evaluate DNA damage accumulated during spaceflight from RNA-Seq data.

### Variant Identification

#### Identifying likely inherited mutations to create an experiment-specific Panel of Normals (PONs)

To examine mutations caused by spaceflight conditions, variants that were likely inherited first needed to be identified and filtered out. We selected a GeneLab transcriptome study of Arabidopsis seedlings grown on the ISS in microgravity or simulated gravity conditions with a paired ground control sample (GLDS-223)^25^. By identifying variants common to spaceflight and ground control vs. ones unique to spaceflight, we could distinguish inherited mutants common to the source seeds from *de novo* variants. The seedlings used in this analysis were grown for five days on the ISS. Root and shoot samples were collected separately. There were two genotypes, wild-type Columbia-0 plants and a transgenic genotype with mammalian type I inositol polyphosphate 5-phosphatase (InsP 5-ptase)^37^. These *Arabidopsis* genotypes are highly inbred and offer a low level of heterozygosity. Shared variants common to each tissue type and Arabidopsis genotype were used to create four Ultimate PONs. The four distinct Ultimate PONs help ensure that inherited mutations shared among the tissue types or genotypes are accounted for when analyzing those samples. These Ultimate PONs are later used as the “known list of population variants” for BaseRecalibrator and the “PON” for Mutect2. HaplotypeCaller consistently called more variants than Mutect2, and the final Ultimate PONs had an average of 7346.75 variants (Table 1).

**Table 1.** Creating the Ultimate PON. An Ultimate Panel of Normals (PON) was created by combining variants called by Mutect2 and HaplotypeCaller. This identifies likely inherited mutants conserved across samples. Mutect2 was run with parameters and steps that would identify mutations shared among the samples. HaplotypeCaller was run in joint genotyping mode to identify shared germline mutations. The Ultimate PON was created using variants called by either tool in at least three of the ten samples per tissue type and *A. thaliana* genotype combination.

|  | Number of Likely Shared Variants from Mutect2 | Number of Joint Genotyped Variants from HaplotypeCaller | Number of Variants in the Ultimate PON |
| --- | --- | --- | --- |
| *Transgenic Shoot* | 10933 | 13291 | 7469 |
| *Transgenic Root* | 6835 | 9505 | 6954 |
| *Wild-Type Shoot* | 11408 | 13902 | 8021 |
| *Wild-Type Root* | 6818 | 9510 | 6943 |

#### Somatic Variant Calling in Arabidopsis

Once the common variants across all samples were removed, the remaining variants were evaluated to assess the impact of spaceflight at the nucleotide level. Each of the 40 samples went through the variant calling and filtering pipeline outlined in Knight et al. 2022^32-33^. Initially, an average of 31,942.08 variants were called by Mutect2. The hard filtering and deduplication steps filtered out the largest number of variants. Hard filters removed 18,349 variants on average, and deduplication filtered an average of 11,899.75 variants. Filtering variants from difficult-to-sequence reads and based on gene expression had less impact on final variant counts, with an average of 605.98 and 15.93 variants filtered out, respectively (Table 2). Final variant counts from each sample’s final VCF were then used for further analysis (Supplementary Table 1).

**Table 2.**
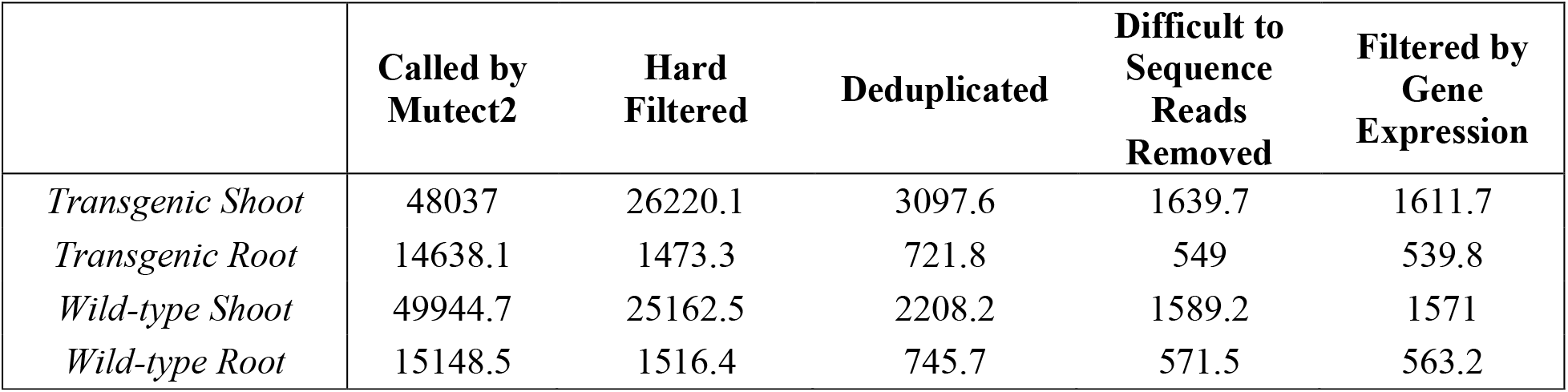
Average number of variants over the filtering steps. The total number of variants called by Mutect2 was filtered for each of the sets of *A. thaliana* samples. Hard filters and deduplication filtered out the largest number of variants, while removing difficult to sequence reads and filtering by gene expression filtered out fewer variants. Shoot tissue samples consistently had a larger number of variants on average compared to root tissue samples, independent of the *A. thaliana* genotype.

|  | Called by Mutect2 | Hard Filtered | Deduplicated | Difficult to Sequence Reads Removed | Filtered by Gene Expression |
| --- | --- | --- | --- | --- | --- |
| *Transgenic Shoot* | 48037 | 26220.1 | 3097.6 | 1639.7 | 1611.7 |
| *Transgenic Root* | 14638.1 | 1473.3 | 721.8 | 549 | 539.8 |
| *Wild-type Shoot* | 49944.7 | 25162.5 | 2208.2 | 1589.2 | 1571 |
| *Wild-type Root* | 15148.5 | 1516.4 | 745.7 | 571.5 | 563.2 |

### Evaluation of the identified variants

#### Spaceflight condition influenced the number of variants called in Arabidopsis

To determine if RNA data from plants grown in space could reveal increased mutation levels compared to ground control, we investigated for differences in the number of mutations identified between spaceflight and. ground control samples. Spaceflight samples were expected to be associated with more mutations than ground control samples because of the increased radiation exposure. This hypothesis was based on the expectation that space radiation greatly affects the mutation rate, as previous research has shown that radiation exposure results in DNA damage^4,22-23^. To investigate this, we compared sample data from three different treatment conditions: ground control, spaceflight in a gravity control (simulated gravity and non-microgravity spaceflight stressors, including space radiation), and normal spaceflight conditions. Spaceflight samples were exposed to a maximum total radiation dose of 5.46 milligrays. The spaceflight group grown in the gravity control had normal spaceflight conditions, except they were centrifuged to simulate gravity. In the original experiment, CO2 conditions and temperature on the ISS were mimicked in the ground control samples to limit the impact of these conditions. For simplicity, we refer to the stress affecting samples grown in the gravity control in space as space radiation. However, we acknowledge that other non-microgravity spaceflight stress conditions could impact this group^4,38-40^. The samples consisted of two tissue types (root and shoot tissue) and two genotypes of Arabidopsis (a wild-type and a transgenic line). The final variant counts were analyzed with a generalized linear model to examine the impact of each spaceflight condition, tissue type, genotype, and any possible interactions (Supplementary Tables 2 & 3).

Interaction effects were investigated first to see if relationships between tissue type, genotype, or spaceflight condition (space radiation and gravity factor) impacted the number of variants called. This is important because any significant variable interactions must be accounted for to interpret results appropriately. Each two-, three-, and four-way interaction was tested for an impact on the number of variants called. Four interaction effects were found to significantly affect the number of variants called (Table 3). The first was a three-way interaction between tissue type, gravity, and radiation. This means all three conditions must be included in statistical analysis to properly determine the effect of tissue type, gravity, and space radiation. The other significant interactions were two-way interactions nested between these three conditions. Thus, the number of variants called in samples exposed to space radiation was analyzed in the context of gravity and tissue type to assess each treatment’s impact properly.

**Table 3.**
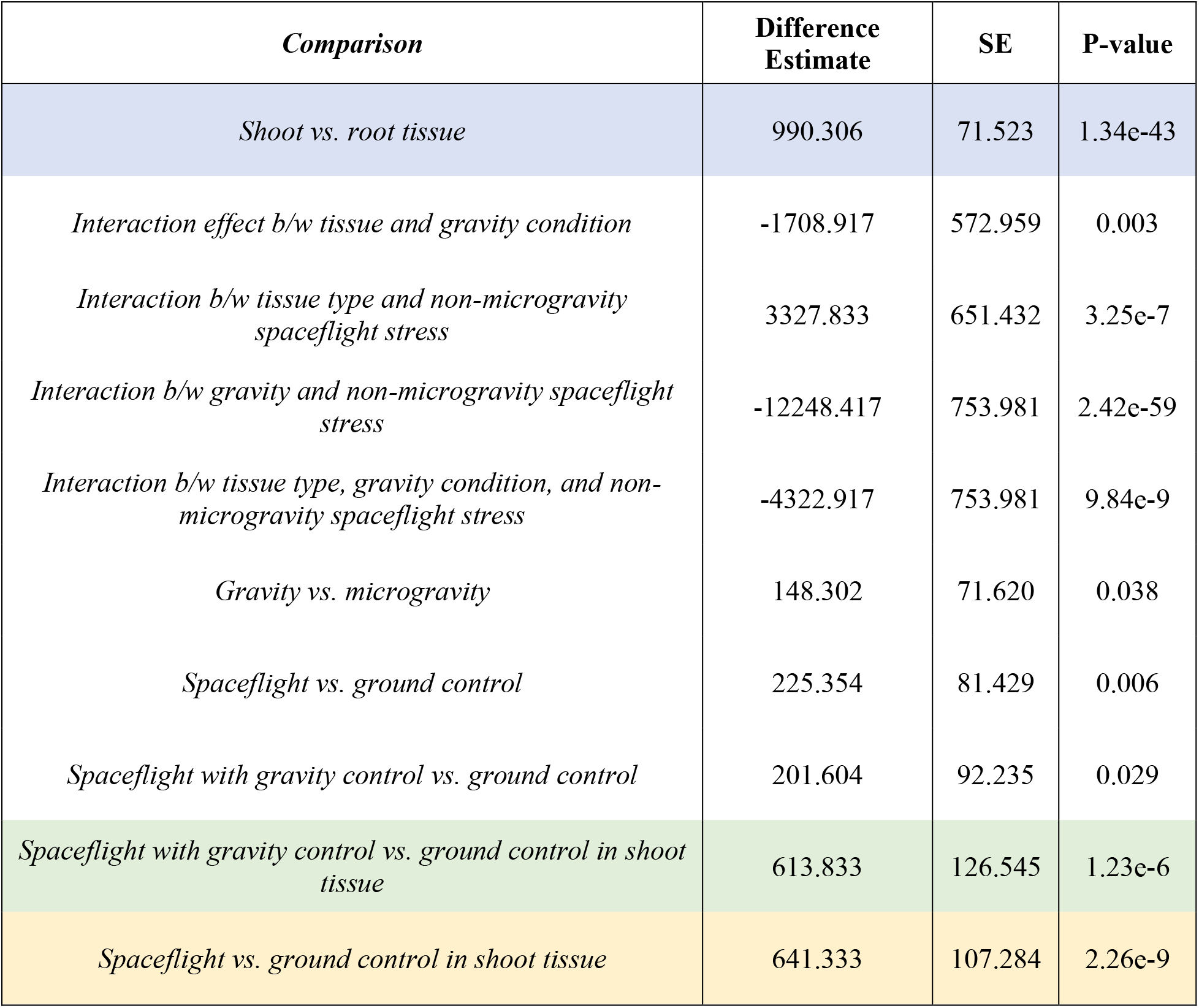
Significant estimates found when comparing treatment groups. This study found that spaceflight is significantly linked to a higher number of variants called in *A. thaliana* samples. Interaction effects were found between the spaceflight conditions and tissue type. When the significant interacting conditions (gravity level, tissue type, and non-microgravity spaceflight stress (namely space radiation)) are incorporated (highlighted in green) it leads to an estimated 613.833 more variants found in shoot tissue grown in a centrifuge on the ISS compared to the ground control. Other interesting estimates include all spaceflight shoot tissue samples vs. ground control tissue samples (highlighted in yellow) and the large difference in the number of variants called between tissue type across all other treatment conditions (highlighted in blue).

Arabidopsis genotype did not impact the number of variants identified (estimate: 19.39, p-value = 0.7863) (Supplementary Table S3, Supplementary Fig. S5, Fig. 2). This was an expected result. The two genotypes of Arabidopsis used in this study were wild-type Col-0 and a transgenic genotype overexpressing the mammalian type I InsP 5-ptase. Since InsP 5-ptase is not an Arabidopsis gene^37^, it did not map to the Arabidopsis reference genome during the read alignment step. Therefore, any reads corresponding to that gene were not considered in downstream analysis. Tissue type heavily influenced spaceflight’s impact on the number of variants called (Supplementary Fig. S5). Without taking individual spaceflight conditions into account, root tissue samples had far fewer variants called compared to shoot tissue samples (estimate: 990.3 variants, p-value = 1.34e-43). The two tissue types were separated for analysis due to the three-way significant interaction effect found between tissue type, gravity, and non-microgravity spaceflight stress. Root tissue did not have any significant differences among the spaceflight conditions (root samples in spaceflight with simulated gravity vs. root samples in normal spaceflight conditions -- estimate: 39.99, p-value=0.72; root samples in spaceflight with simulated gravity vs. root samples grown on Earth -- estimate: 210.63, p-value = 0.12) (Supplementary Table S3).

**Figure 2.**
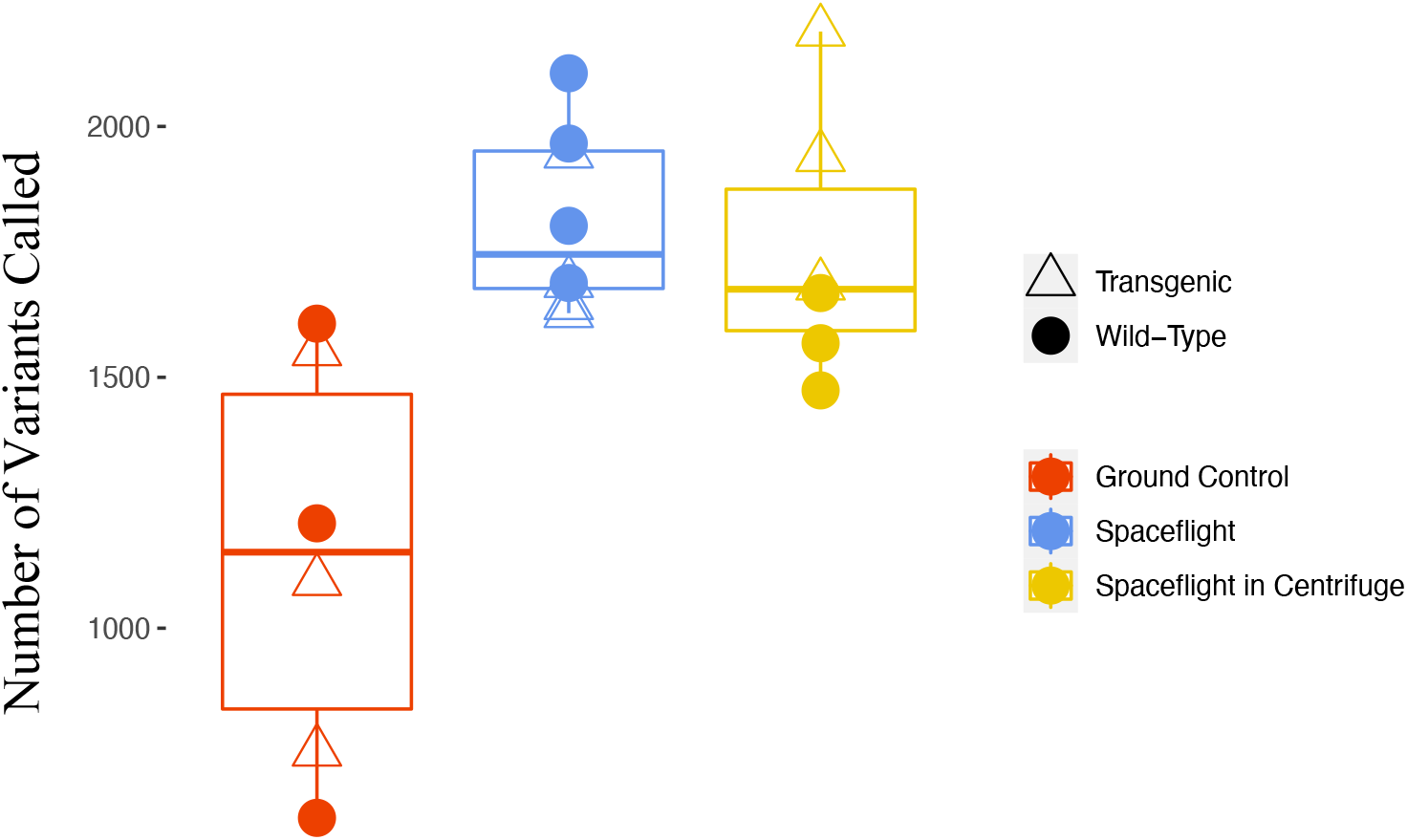
Variant counts in the shoot tissue *A. thaliana* samples. The number of variants found (y-axis) for each shoot tissue sample. Both sets of spaceflight (yellow and blue) had a higher number of variants called compared to the ground control samples (red). The samples grown in space in a gravity control (yellow) also had a higher number of variants compared to the ground control samples (red). The spaceflight group in a gravity control allows for a focus on the non-microgravity spaceflight stress conditions, particularly space radiation, without exposure to microgravity.

The number of variants found in the shoot tissue, on the other hand, was significantly different among the spaceflight conditions (Fig. 2). Overall, shoot tissue samples grown aboard the ISS had a higher number of variants compared to the ground control samples (estimate: 641.33, p-value = 2.26e-09). However, the impact of the spaceflight conditions needed to be separated out because of the significant interaction effect observed between tissue type, gravity, and non-microgravity spaceflight stress, such as space radiation. Shoot tissue samples grown in simulated gravity aboard the ISS had more mutations than the ground control samples (estimate: 613.83, p-value = 1.23e-06). This comparison points to space radiation, rather than microgravity, as the factor that induces more DNA damage. Unfortunately, there were no samples grown in microgravity that were not also exposed to space radiation which would have allowed for a closer examination of space radiation’s impact at the nucleotide level. However, the impact of gravity and microgravity was investigated by comparing the two groups of samples grown on the ISS. These ISS-grown samples were all exposed to space radiation but were grown in either microgravity or simulated gravity conditions. The impact of microgravity was not significant in the shoot tissue samples grown aboard the ISS (estimate: 55.00, p-value = 0.642). This supports a link between space radiation and an accumulation of variants; however, an experiment examining simulated microgravity vs. normal gravity conditions on Earth would add to our understanding of microgravity’s role in the mutation rate.

#### Spaceflight influenced the types of variants identified Arabidopsis

Radiation has been linked to a higher number of transversion mutations, motivating us to examine whether a higher number of transversion mutations could be observed in the spaceflight samples selected for this study^22-23^. Each sample’s VCF was analyzed using a custom script to examine the types of mutations identified. The ratio of transversion mutations to the total number of SNPs was analyzed using a generalized linear model with a binomial distribution, with the ratio as a proportion. The model’s predictor variables were spaceflight conditions, Arabidopsis genotype, and tissue type. These variables were investigated to determine their impact on the proportion of transversion mutations (Supplementary Fig. S6). Once again, the Arabidopsis genotype did not significantly impact the proportion of transversion mutations (estimate: 0.008, p-value: 0.212). Interactions between tissue type, gravity, and space radiation level and all nested two-way interactions were also significant (Table 4). Thus, all three conditions were incorporated into the model to investigate the impact of spaceflight.

**Table 4.**
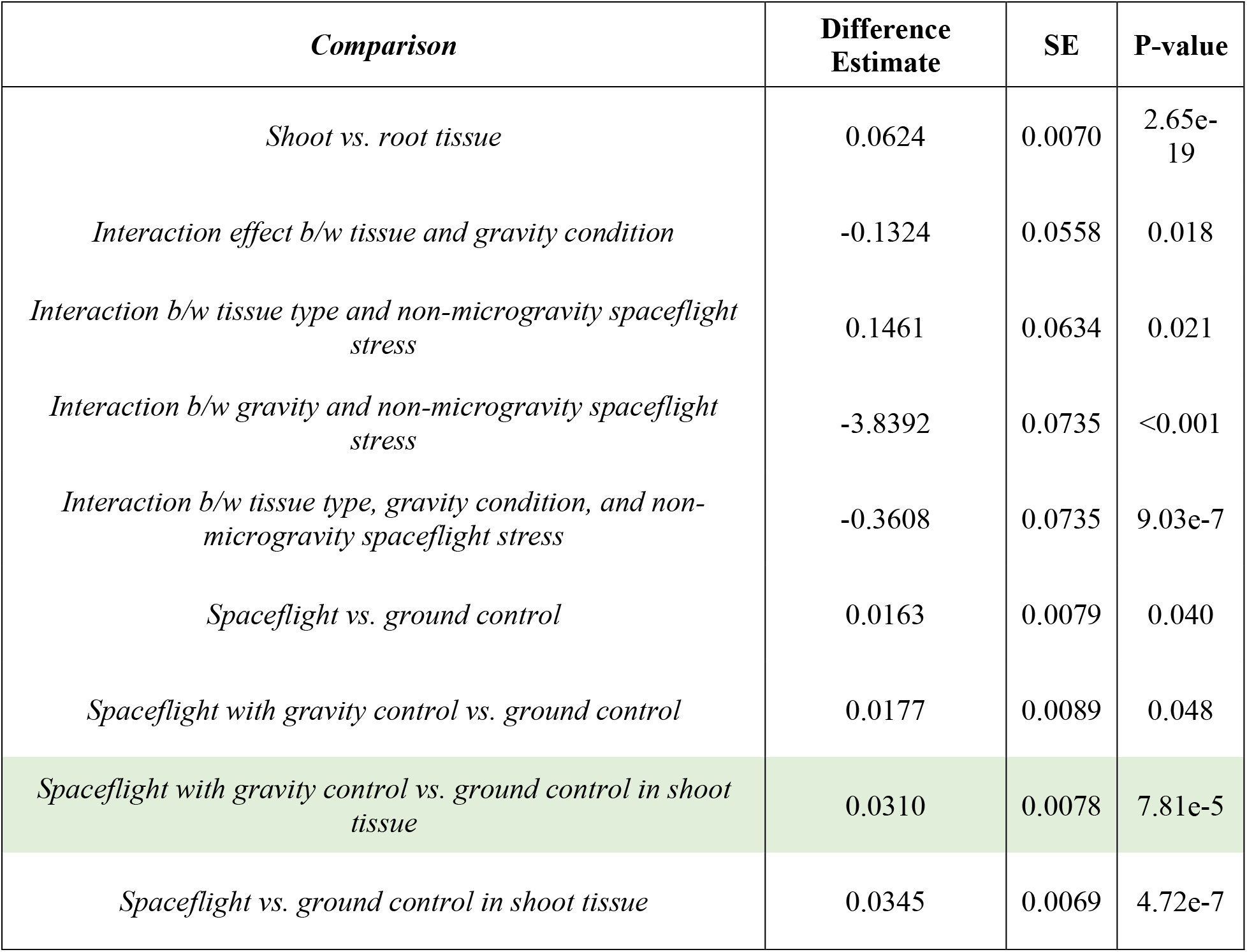
Significant differences in the proportion of transversion mutations found when comparing treatment groups. Spaceflight significantly impacts the proportion of transversion mutations found. Interaction effects were found between the spaceflight conditions and tissue type. When the significant interactions (gravity level, tissue type, and non-microgravity spaceflight stress (namely space radiation)) are incorporated (highlighted in green) it leads to an estimated increase of 0.031 in proportion of transversion mutations. This was found in shoot tissue grown in a centrifuge on the ISS compared to the ground control.

| *Comparison* | **Difference Estimate** | **SE** | **P-value** |
| --- | --- | --- | --- |
| *Shoot vs. root tissue* | 0.0624 | 0.0070 | 2.65e-19 |
| *Interaction effect b/w tissue and gravity condition* | -0.1324 | 0.0558 | 0.018 |
| *Interaction b/w tissue type and non-microgravity spaceflight stress* | 0.1461 | 0.0634 | 0.021 |
| *Interaction b/w gravity and non-microgravity spaceflight stress* | -3.8392 | 0.0735 | <0.001 |
| *Interaction b/w tissue type, gravity condition, and non-microgravity spaceflight stress* | -0.3608 | 0.0735 | 9.03e-7 |
| *Spaceflight vs. ground control* | 0.0163 | 0.0079 | 0.040 |
| *Spaceflight with gravity control vs. ground control* | 0.0177 | 0.0089 | 0.048 |
| *Spaceflight with gravity control vs. ground control in shoot tissue* | 0.0310 | 0.0078 | 7.81e-5 |
| *Spaceflight vs. ground control in shoot tissue* | 0.0345 | 0.0069 | 4.72e-7 |

Spaceflight shoot tissue samples had a higher proportion of transversion mutations compared to the ground control shoot tissue samples (estimate: 0.035, p-value = 4.72e-07) (Table 4, Fig. 3). Due to the significant interaction effects found, samples from each spaceflight group were analyzed separately to draw appropriate conclusions. Spaceflight shoot tissue samples grown in simulated gravity aboard the ISS had a larger proportion of transversion mutations compared to shoot tissue samples grown on the ground (estimate: 0.031, p-value 7.81e-05) (Table 4, Fig. 3). This comparison allows a closer examination of the impact space radiation has on the proportion of transversion mutations identified and is further evidence that radiation leads to a higher rate of transversion mutations, as demonstrated in a few previous studies^22-23^. Root tissue samples did not have significant differences in the proportion of transversion mutations among either spaceflight condition, which is consistent with the reduced number of all single nucleotide variants identified in root tissue.

**Figure 3.**
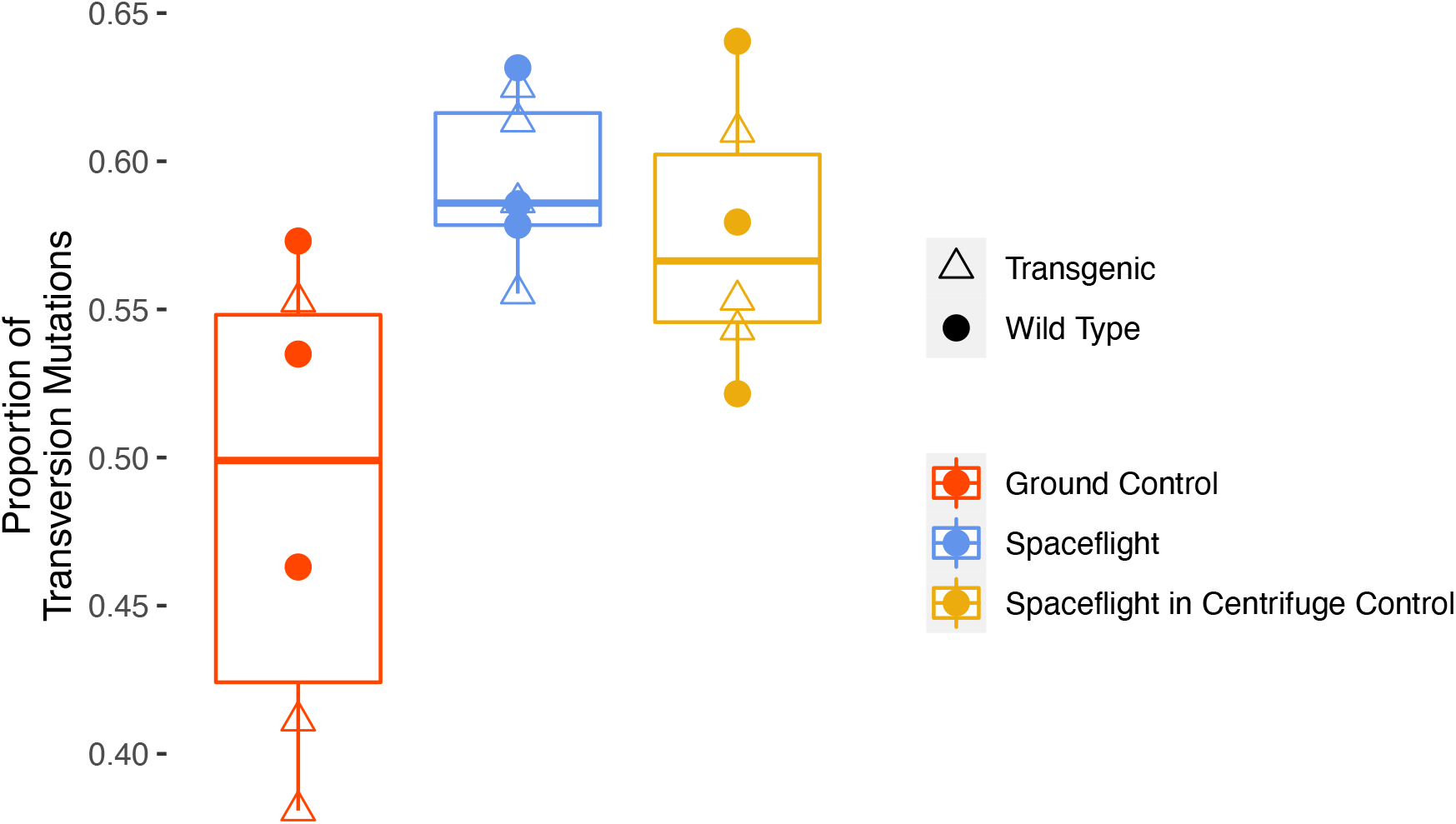
Proportion of transversion mutations identified in the shoot tissue *A. thaliana* samples. The spaceflight groups (yellow and blue) had a higher proportion of transversion mutations compared to the ground control samples (red). Samples in the gravity control that were exposed to space radiation (yellow) also had a higher proportion of transversions compared to the ground control samples (red), demonstrating that non-microgravity spaceflight stress, like space radiation, is a main contributor to the higher proportion of transversion mutations.

We also examined the types of transversion point mutations enriched in spaceflight (Table 5). G → C/C → G mutations and G → T/C→ A mutations were significantly enriched in the spaceflight shoot samples grown in simulated gravity (Table 5). The other two types of transversion mutations, A → C/T → G and A → T/T → A, were not enriched in the spaceflight samples. Overall, our approach allowed us to identify nucleotide composition changes. We observed increased variation during spaceflight, which likely indicates DNA damage from radiation. Based on the centrifuge control, microgravity does not appear to impact the number of variants detected in this study.

**Table 5.**
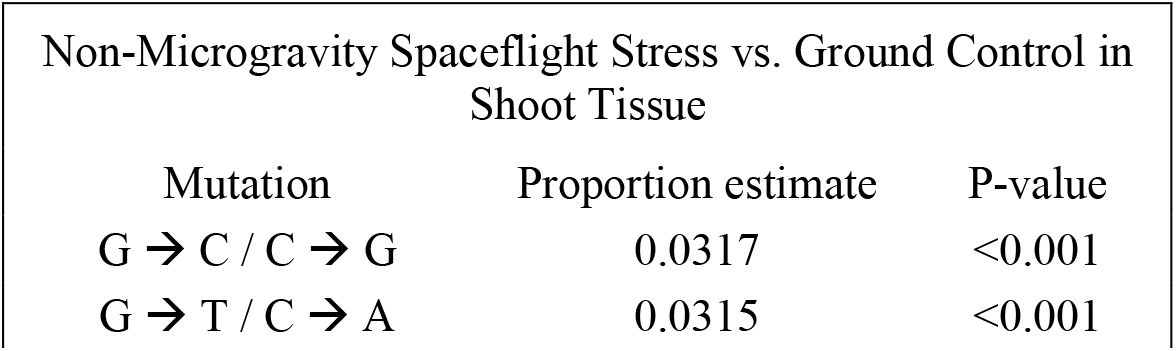
Specific transversion point mutations enriched in spaceflight conditions. Shoot tissue remained the tissue type of interest when examining types of point mutations. Two types of transversion point mutations were significantly enriched in spaceflight when comparing samples grown with the gravity control in space to the ground control samples. This allowed for a closer examination of the effects of spaceflight without the influence of microgravity.

| Non-Microgravity Spaceflight Stress vs. Ground Control in Shoot Tissue |  |  |
| --- | --- | --- |
| Mutation | Proportion estimate | P-value |
| G → C / C → G | 0.0317 | <0.001 |
| G → T / C → A | 0.0315 | <0.001 |

## Discussion

Space travel presents new challenges to terrestrial life, including high levels of space radiation and microgravity. Studying the effects of spaceflight on terrestrial life in true spaceflight conditions is the best way to study its impact, but this is challenging due to costs and access. This study took advantage of publicly available RNA-Seq data from NASA’s GeneLab to identify variants in Arabidopsis samples grown aboard the ISS. The number of variants identified per sample was investigated to examine the impact of spaceflight at the nucleotide level. Even though the plants were only grown for five days on the ISS, we were able to detect an increase in variants in Arabidopsis samples grown in spaceflight.

Examining the specific impact of spaceflight stress is difficult since multiple stressors are occurring simultaneously. Because of the unique experimental design of the GeneLab data set, we were able to examine the impact of non-microgravity spaceflight stress such as space radiation. This was done by comparing Arabidopsis samples grown in simulated gravity aboard the ISS to samples under ground control conditions. Spaceflight stress led to a significantly higher number of called variants than the ground control for shoot tissue samples (estimate: 613.83, p-value = 1.23e-06). However, we did not observe a statistical difference between the spaceflight samples in microgravity and in the centrifuge (root estimate: 39.99, p-value = 0.72; shoot estimate: 55, p-value =0.64). This suggests that microgravity may not affect the number of nucleotide changes occurring. This is surprising as there are reports that microgravity affects relevant pathways for mitigating damage, including effects on DNA repair enzymes and impacts on cellular proliferation observed in astronauts^4-7^. The lack of effect observed due to microgravity in this analysis could be due to the short time samples were on the ISS. Future studies could examine microgravity’s impact over longer periods of time to better understand its effect on DNA. A limitation of this analysis was the lack of samples grown on Earth in simulated microgravity. This would have allowed for microgavity’s impact to be better understood. Future experiments which examine microgravity and radiation both separately and together would be extremely useful for reflecting true spaceflight conditions.

A significant interaction effect was observed between tissue type, gravity, and space radiation level. This meant there was an observable relationship between these three conditions. To properly determine the impact of any of the treatments required the incorporation of the other two conditions as well. Spaceflight conditions were significantly associated with a higher number of variants in the shoot tissue samples. However, no observable trend was detected in root tissue. The interaction was broken down into the two spaceflight groups: spaceflight with and without the gravity control. In the shoot tissue samples, both spaceflight groups were associated with a higher number of variants. However, in the root tissue, there was no significant difference in the number of variants called in the spaceflight groups compared to the ground control samples. This difference between shoot and root tissue resulted in the three-way interaction effect between tissue type, gravity, and non-microgravity spaceflight stress that was included in the statistical analysis.

One potential reason for the significant interaction effect could be issues with the number of variants called in the root tissue samples. This group was associated with a higher standard error than any other treatment group (Supplementary Table S2). The uncertainty in the number of variants called in the root tissue samples could be due to a few things. For one, the transcript composition in the root and shoot tissue varied greatly (Supplementary Figure S4). Shoot tissue samples had more highly expressed genes compared to the root tissue samples which could have led to a higher number of variant calls. We observe these differences in transcript composition in other RNA-seq data from Arabidopsis. This difference in expression points out a major challenge of calling variants from RNA-seq data; that it is dependent on the transcript expression level. Here, the expression level per gene differs significantly between root and shoot and it affects the ability to identify variants in the root tissue. Another issue is the sample size, there were only two ground control root tissue samples per Arabidopsis genotype. This could contribute to the high standard errors for this group, which would influence the statistical analysis. Future experiments could increase the number of samples per treatment group to decrease standard error and potentially find significant treatment effects. Determining if the tissue specific differences we observe are due to actual biological differences will likely require a direct analysis of the DNA or approaches to control for differences in expression between root and shoot tissues. However, this is a critical question to answer as it could provide insights into tissue-specific DNA repair differences or radiation sensitivity..

Variant composition was examined to investigate the types of variants accumulating in space and allow for a closer examination of the impact spaceflight had at the nucleotide level. The results showed transversion mutations were more common in spaceflight, with space radiation believed to be the contributing treatment. Transversion mutations have previously been shown to be more common in the presence of radiation^22-23^. The proportion of G → T / C → A transversions was significantly higher in shoot tissue samples in the simulated gravity that were exposed to space radiation vs. the ground control samples that were not exposed to space radiation. This aligns with previous findings that linked UVA radiation from the sun to a higher proportion of these sorts of transversion^23^. Additionally, the proportion of C → G / G → C transversions increased when examining the effect of space radiation on shoot tissue samples grown in simulated gravity (Table 5). This is also consistent with an increased risk of these transversions due to exposure to solar radiation22. The difference in the proportions of the other types of point mutations may have also been significant if the samples were in spaceflight for longer than five days. A study examined mutations accumulated in astronauts using DNA^11^. Even though they were only aboard the Space Station for a median of 12 days, a significant number of mutations was observed^11^. In contrast to our study, they observed a higher number of transition mutations. These differences could be due to the smaller number of genes they examined, differences between human and Arabidopsis sensitivity to DNA damage and repair, or differences in comparing mutations identified from RNA versus directly from DNA. Nevertheless, it does further demonstrate that more research is needed to understand DNA damage caused by spaceflight. Duration of exposure, impact on different species or tissue type, and individual spaceflight conditions could all be further studied to better understand spaceflight’s impact on DNA.

NASA’s new ‘Omics database, GeneLab, provides the opportunity to examine results from spaceflight related experiments without having to carry out an additional investigation. This is a tremendous resource for the scientific community because experiments on the ISS are difficult to secure and extremely expensive. Nonetheless, most of the information on GeneLab is related to gene expression. Both *Mus musculus*, which comprises almost a third of the GeneLab datasets and Arabidopsis, which makes up the significant majority of the plant data, are highly inbred species; we specifically tailored our pipeline to identify variants in inbred species. This study demonstrated DNA damage can be investigated using sequencing information from RNA-Seq experiments. This is advantageous for examining the impact spaceflight-related stress as there is a large amount of RNA-Seq data on GeneLab that could be examined for this purpose. This analysis could also be applied to investigate damage caused by other stress conditions. However, one limitation in this study is that the called variants could be the result of spaceflight-specific RNA-editing events and not DNA damage. This is not something we are able to separate out. Samples that directly compare DNA damage from space radiation with associated RNA-Seq would allow for a complete evaluation of this approach. This would clarify if the differences we observe are due to spaceflight-based changes directly on the DNA or RNA processing events.

## Conclusions

This study used the high sequence resolution of RNA-Seq data to examine mutation rate caused by spaceflight stress. Publicly available RNA-Seq data was downloaded from NASA’s GeneLab and investigated for an accumulation of variants in spaceflight conditions. There were three treatment conditions examined: ground control, spaceflight with a gravity control, and normal spaceflight conditions. Space radiation contributed to a higher number of detected variants in shoot tissue samples. A higher number of variants were found in the shoot tissue samples grown on the ISS compared to the ground control shoot tissue samples, but this trend was not observed in root tissue. This study also linked spaceflight stress to a higher number of transversion mutations in shoot tissue samples. Long-term space travel is becoming more of a reality and observing how spaceflight stress results in nucleotide changes can help to understand and combat the risks. Here we show that the vast resource of RNA-Seq data available from spaceflight samples can be used to identify nucleotide damage.

## Acknowledgements

This work was supported by a grant from the National Aeronautics and Space Administration (80NSSC19K0146) to Dr. Colleen J. Doherty. This work was also supported in part by NIEHS training grant T32ES007329 (M.K.). We thank Dr. Jason A. Osborne from the Department of Statistics at North Carolina State University for guidance on appropriate statistical analysis techniques. We also thank Dr. Imara Y. Perera from the Department of Plant and Microbial Biology at North Carolina State University for feedback, review, and excitement regarding this analysis and manuscript.

