## Supplementary Information for "Assessing the Nucleotide-Level Impact of Spaceflight Stress using RNA-Sequencing Data"

### Supplemental Figures

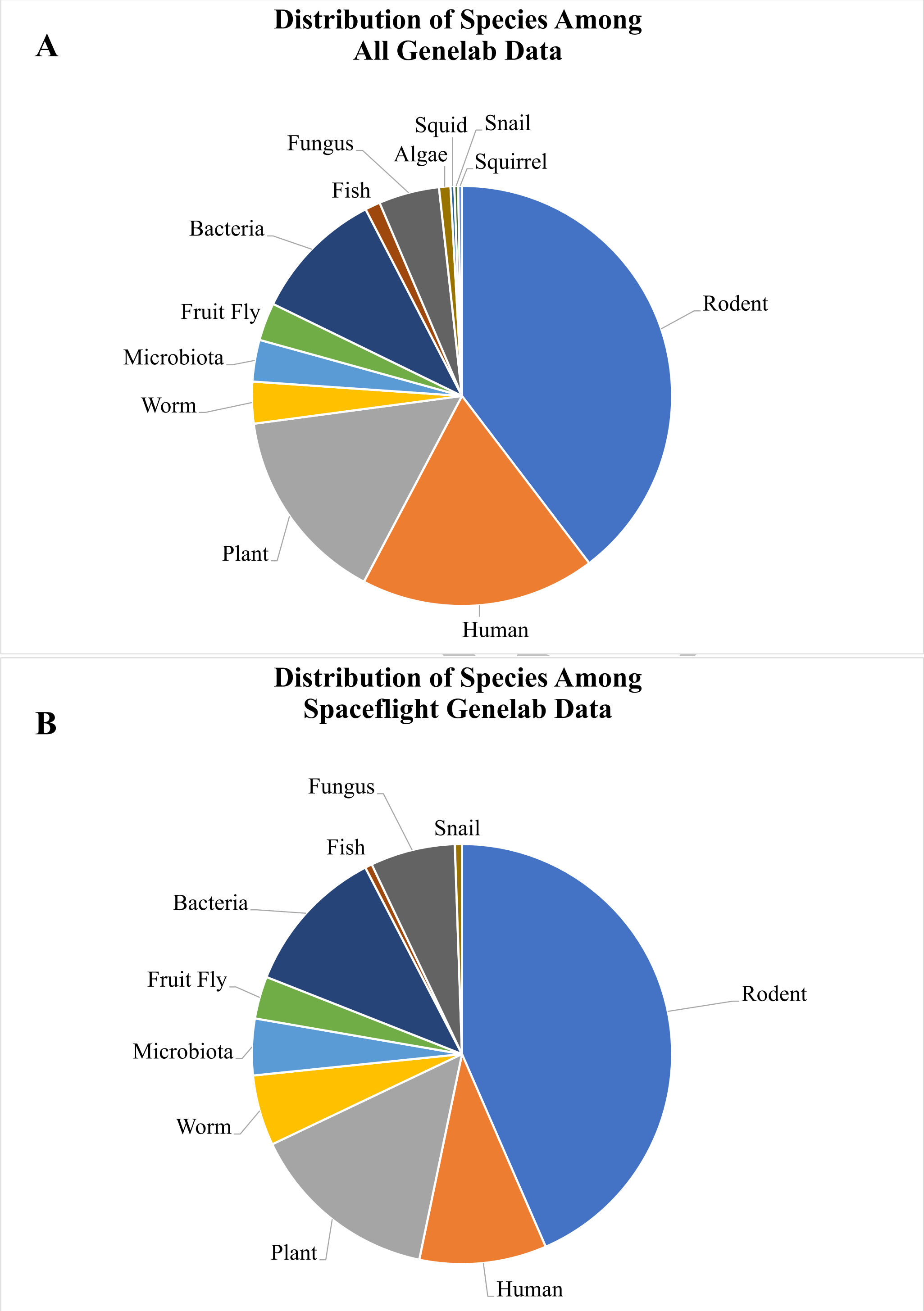


**Supplementary Figure S1. Types of species with data available on GeneLab.** (Data Accessed August 29, 2022). There are many different types of species with data available on GeneLab. (A) Rodents are the most common species with data in all the GeneLab data and (B) the spaceflight centric GeneLab data. Plant data encompasses many experiments on GeneLab and is the second most common type of data in the spaceflight experiments.

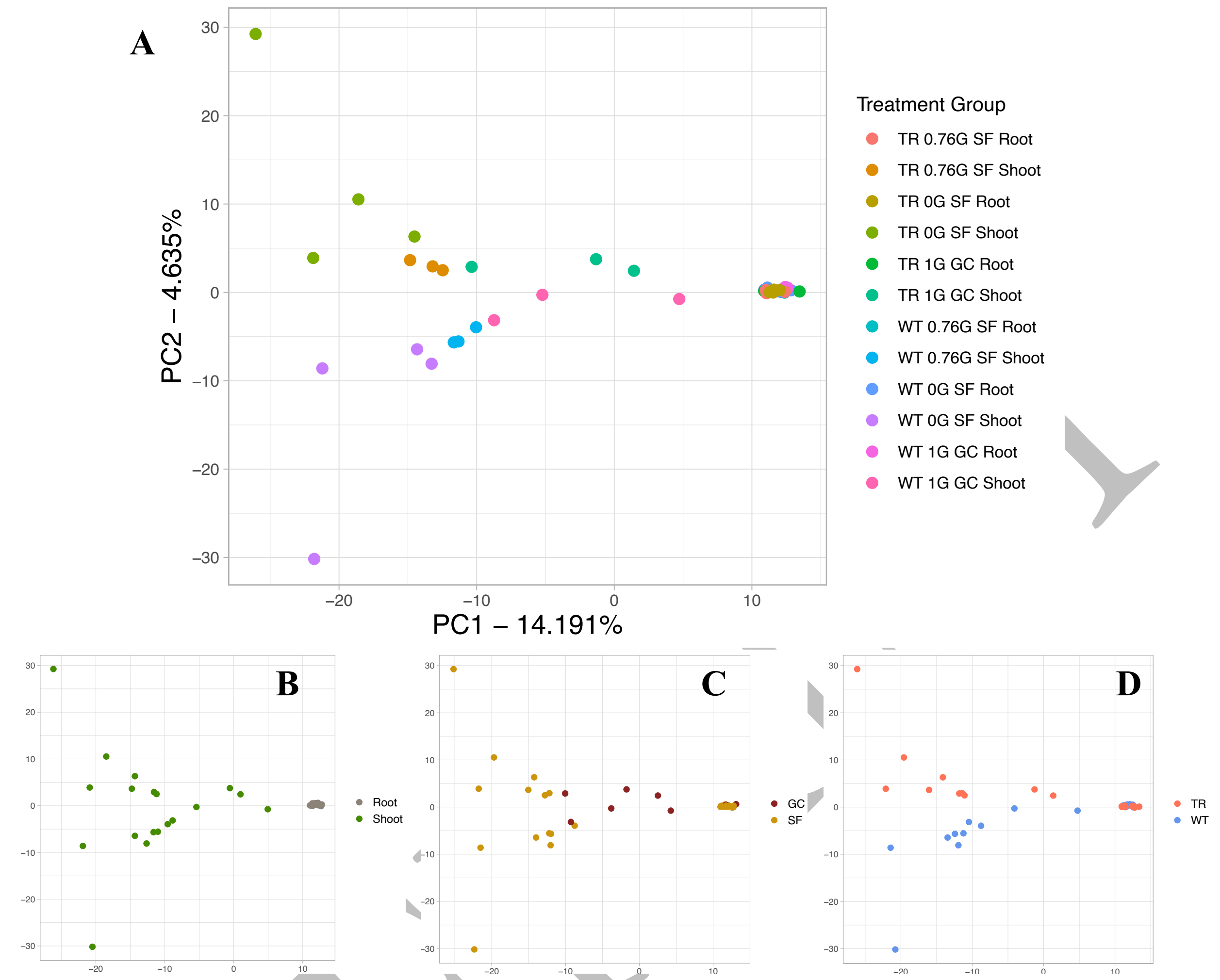


**Supplementary Figure S2. Variant counts per gene principal component analysis.** The x-axis shows the variance across Principal Component 1 (PC1). PC1 explained 14.19% of the variance. The y-axis shows Principal Component 2 (PC2) which explained 4.64% of the variance. (A) The variance explained by the first two principal components separates the treatment combinations of *A. thaliana*. Tissue type is the main contributor to the first principal component (B), with spaceflight and ground control conditions also explaining some of the variation (C). *A. thaliana* genotype seems to explain the variation on the second principal component (D).

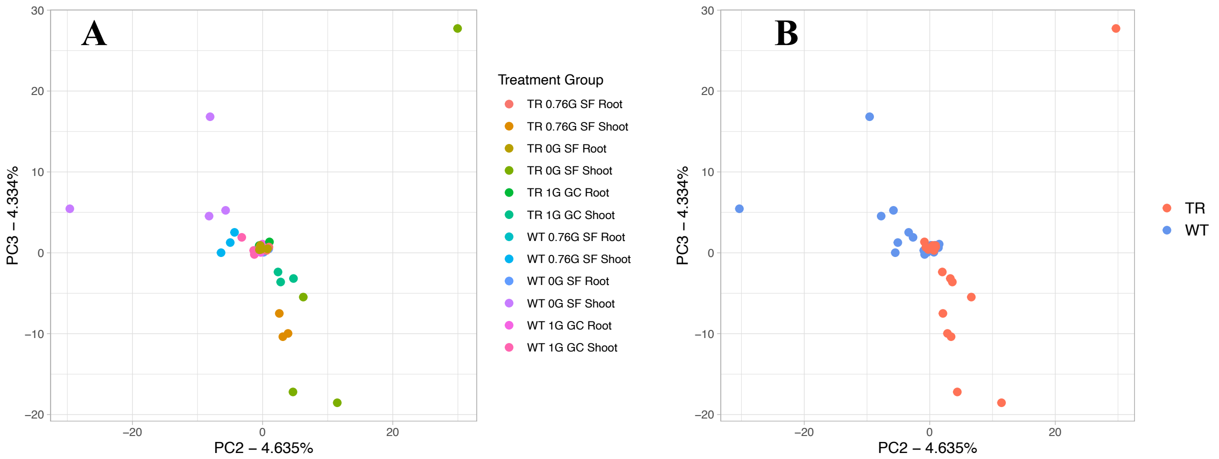


**Supplementary Figure S3. Variant counts per gene principal components 2 vs. 3.** The x-axis shows the variance across Principal Component 2 (PC2). PC2 explained 4.64% of the variance. The y-axis shows Principal Component 3 (PC3) which explained 4.33% of the variance. The variance described by second and third principal components explains less of the variance among the treatment combinations of *A. thaliana* (A). It does appear that *A. thaliana* genotype is the main contributor to variation in both the second and third principal component (B).

PREPRINT

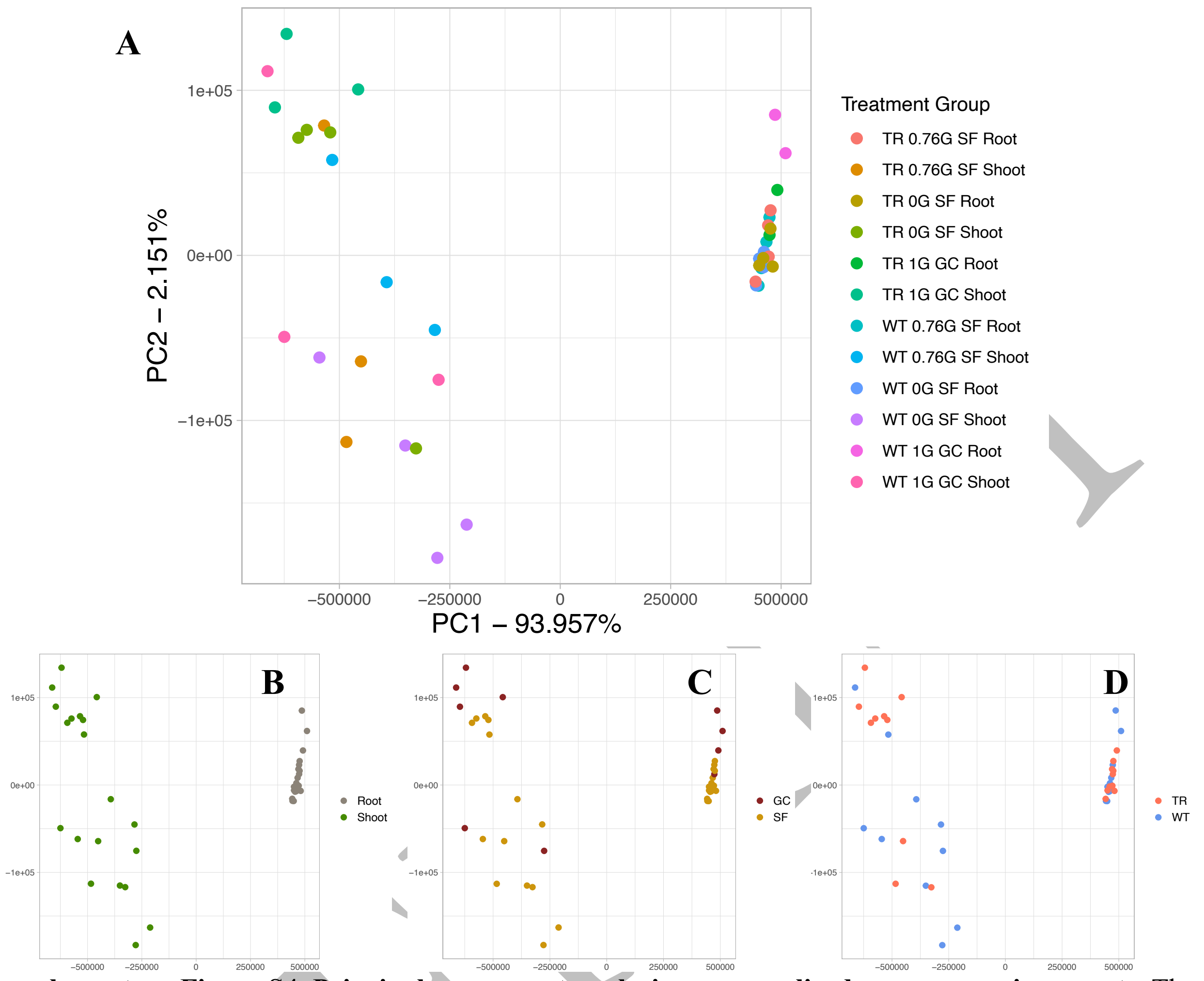


**Supplementary Figure S4. Principal component analysis on normalized gene expression counts.** The x-axis shows the variance across Principal Component 1 (PC1). PC1 explained 93.96% of the variance. The y-axis shows Principal Component 2 (PC2) which explained 2.15% of the variance. The variance explained by the first two principal components does not group the treatment combinations of *A. thaliana* (A). Tissue type is clearly accounting for the variance in the first principal component (B). Spaceflight condition (C) and *A. thaliana* genotype do not have as much impact on variance explained by either the first or second principal component (D).

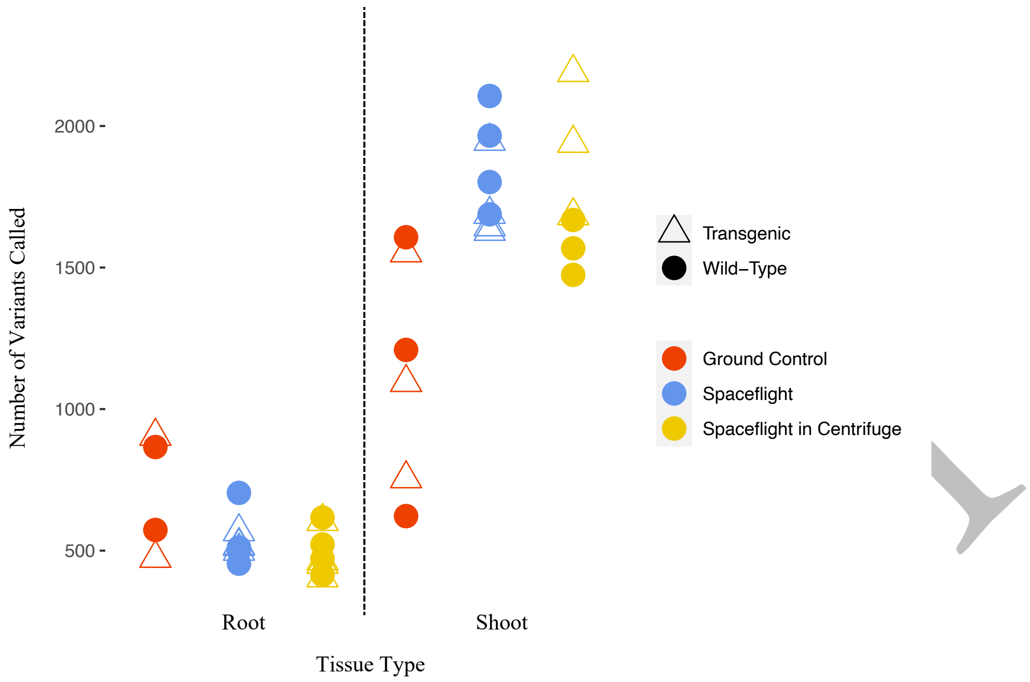


**Supplementary Figure S5. Variant counts derived from RNA-Seq data in *A. thaliana* samples.** The number of variants found (y-axis) for each sample. Root tissue samples (left) had a lower number of variants called compared to shoot tissue samples (right). *A. thaliana* genotype (shape) had no significant impact on the number of variants called. In shoot tissue samples (right), the samples grown in spaceflight (yellow and blue) had a higher number of variants called compared to the ground control samples (red).

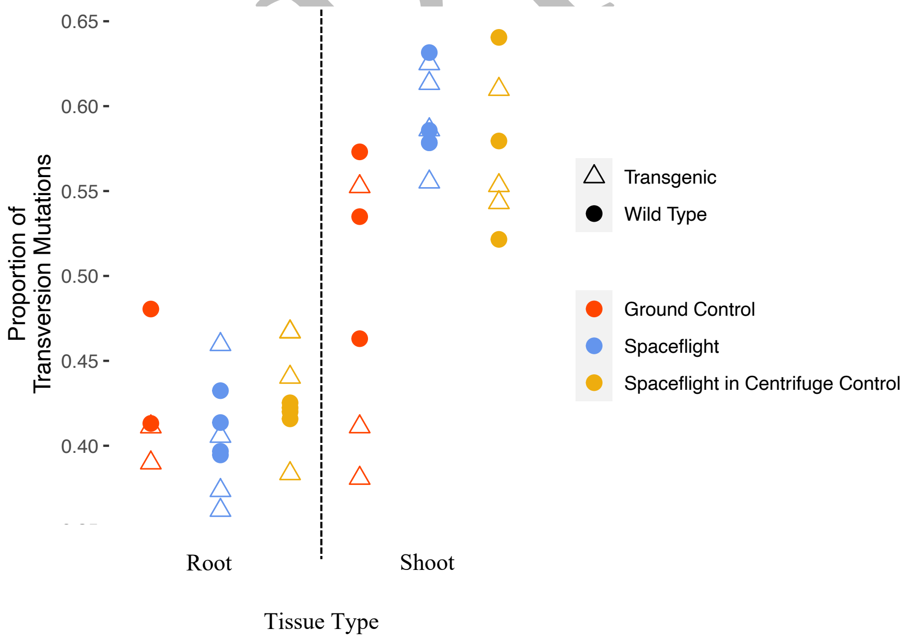


**Supplementary Figure S6. Proportion of transversion mutations identified in RNA-Seq data in *A. thaliana* samples.** The proportion of transversion mutations (y-axis) for each sample. The trends seen are like those found in the overall variant counts of each sample. Shoot tissue samples (right) consistently had a higher number of variants compared to root tissue samples (left). Spaceflight conditions (blue and yellow) were associated with a higher number of variants in the shoot tissue samples (right).

### Supplemental Tables

| *Sample* | Line | Tissue Type | Treatment | Response |
|---|---|---|---|---|
| *FLS18* | Wild-Type | Shoot | Spaceflight with Gravity Control | 1568 |
| *FLS28* | Wild-Type | Shoot | Normal Spaceflight | 1802 |
| *FLS38* | Wild-Type | Shoot | Normal Spaceflight | 1688 |
| *FLS48* | Wild-Type | Shoot | Spaceflight with Gravity Control | 1668 |
| *FLS58* | Wild-Type | Shoot | Spaceflight with Gravity Control | 1474 |
| *FLS68* | Wild-Type | Shoot | Normal Spaceflight | 2106 |
| *FLS78* | Wild-Type | Shoot | Normal Spaceflight | 1966 |
| *GCS1* | Wild-Type | Shoot | Ground Control | 1209 |
| *GCS14* | Wild-Type | Shoot | Ground Control | 1607 |
| *GCS18* | Wild-Type | Shoot | Ground Control | 622 |
| *FL08* | Wild-Type | Root | Spaceflight with Gravity Control | 617 |
| *FL18* | Wild-Type | Root | Spaceflight with Gravity Control | 522 |
| *FL28* | Wild-Type | Root | Normal Spaceflight | 500 |
| *FL38* | Wild-Type | Root | Normal Spaceflight | 511 |
| *FL48* | Wild-Type | Root | Spaceflight with Gravity Control | 414 |
| *FL58* | Wild-Type | Root | Spaceflight with Gravity Control | 471 |
| *FL68* | Wild-Type | Root | Normal Spaceflight | 454 |
| *FL78* | Wild-Type | Root | Normal Spaceflight | 704 |
| *GC14* | Wild-Type | Root | Ground Control | 573 |
| *GC16* | Wild-Type | Root | Ground Control | 866 |
| *GCS13* | Transgenic | Shoot | Ground Control | 1552 |
| *GCS17* | Transgenic | Shoot | Ground Control | 754 |
| *GCS2* | Transgenic | Shoot | Ground Control | 1094 |
| *FLS23* | Transgenic | Shoot | Normal Spaceflight | 1628 |
| *FLS33* | Transgenic | Shoot | Normal Spaceflight | 1644 |
| *FLS63* | Transgenic | Shoot | Normal Spaceflight | 1688 |
| *FLS73* | Transgenic | Shoot | Normal Spaceflight | 1946 |
| *FLS13* | Transgenic | Shoot | Spaceflight with Gravity Control | 1939 |
| *FLS43* | Transgenic | Shoot | Spaceflight with Gravity Control | 1683 |
| *FLS53* | Transgenic | Shoot | Spaceflight with Gravity Control | 2189 |
| *GC01* | Transgenic | Root | Ground Control | 904 |
| *GC13* | Transgenic | Root | Ground Control | 473 |
| *FL23* | Transgenic | Root | Normal Spaceflight | 518 |
| *FL33* | Transgenic | Root | Normal Spaceflight | 515 |
| *FL63* | Transgenic | Root | Normal Spaceflight | 567 |
| *FL73* | Transgenic | Root | Normal Spaceflight | 498 |
| *FL03* | Transgenic | Root | Spaceflight with Gravity Control | 404 |

PREPRINT

| | | | | |
|---|---|---|---|---|
| *FL13* | Transgenic | Root | Spaceflight with Gravity Control | 603 |
| *FL43* | Transgenic | Root | Spaceflight with Gravity Control | 452 |
| *FL53* | Transgenic | Root | Spaceflight with Gravity Control | 464 |

**Supplementary Table S1. Final number of variants called for each sample in the GLDS-223 dataset.** (Leftmost column) Original sample name. (Middle three columns) The corresponding *A. thaliana* genotype, tissue type, and spaceflight treatment group per sample. (Rightmost column) final number of variants called by Mutect2 after all filtering steps.

| Genotype | Tissue Type | Treatment | LSMean Estimate | Standard Error |
|---|---|---|---|---|
| Wild-Type | Shoot | Ground Control | 1146.00 | 126.55 |
| Wild-Type | Shoot | Spaceflight with Gravity Control | 1570.00 | 126.55 |
| Wild-Type | Shoot | Spaceflight | 1890.50 | 109.59 |
| Wild-Type | Root | Ground Control | 719.50 | 154.99 |
| Wild-Type | Root | Spaceflight with Gravity Control | 506.00 | 109.59 |
| Wild-Type | Root | Spaceflight | 542.25 | 109.59 |
| Transgenic | Shoot | Ground Control | 1133.33 | 126.55 |
| Transgenic | Shoot | Spaceflight with Gravity Control | 1937.00 | 126.55 |
| Transgenic | Shoot | Spaceflight | 1726.50 | 109.59 |
| Transgenic | Root | Ground Control | 688.50 | 154.99 |
| Transgenic | Root | Spaceflight with Gravity Control | 480.75 | 109.59 |
| Transgenic | Root | Spaceflight | 524.50 | 109.59 |

**Supplementary Table S2. Least squares estimates for each treatment group.** The least squares estimate for the number of variants per group (*A. thaliana* genotype, tissue type, and spaceflight treatment) as determined by a generalized linear model. Each LSMean estimate represents an estimated average number of variants called in that treatment group. Each estimate has a corresponding standard error that helps to understand the variability among the samples. These estimates are used in the statistical analysis to compare and investigate the treatment impacts.

| *Comparison* | Difference Estimate | SE | P-value |
|---|---|---|---|
| *Shoot vs. root tissue* | -990.306 | 71.523 | 1.34e-43 |
| *Wild-type vs. transgenic genotype* | 19.389 | 71.523 | 0.786 |
| *Interaction effect b/w tissue and genotype* | -264.333 | 429.136 | 0.538 |
| *Interaction effect b/w tissue and gravity condition* | -1708.917 | 572.959 | 0.003 |
| *Interaction b/w tissue type and non-microgravity spaceflight stress* | 3327.833 | 651.432 | 3.25e-7 |
| *Interaction b/w A. thaliana genotype and gravity condition* | -661.583 | 572.959 | 0.248 |
| *Interaction b/w A. thaliana genotype and non-microgravity spaceflight stress* | -247.333 | 651.432 | 0.704 |
| *Interaction b/w gravity and non-microgravity spaceflight stress* | -12248.417 | 753.981 | 2.42e-59 |
| *Interaction b/w tissue type, A. thaliana genotype, and gravity condition* | -703.083 | 572.959 | 0.220 |
| *Interaction b/w tissue type, A. thaliana genotype, and non-microgravity spaceflight stress* | -209.333 | 651.432 | 0.748 |
| *Interaction b/w tissue type, gravity condition, and non-microgravity spaceflight stress* | -4322.917 | 753.981 | 9.84e-9 |
| *Interaction b/w A. thaliana genotype, gravity condition, and non-microgravity spaceflight stress* | -792.583 | 753.981 | 0.293 |
| *Interaction b/w tissue type, A. thaliana genotype, gravity condition, and non-microgravity spaceflight stress* | -648.083 | 753.981 | 0.390 |
| *Gravity vs. microgravity* | 148.302 | 71.620 | 0.038 |
| *Gravity vs. microgravity in space* | 47.500 | 80.657 | 0.556 |
| *Gravity vs. microgravity in space in shoot tissue* | 55.000 | 118.372 | 0.642 |
| *Gravity vs. microgravity in space in root tissue* | 40.000 | 109.592 | 0.715 |

PREPRINT

| | | | |
|---|---|---|---|
| *Spaceflight vs. ground control* | 225.354 | 81.429 | 0.006 |
| *Spaceflight with gravity control vs. ground control* | -201.604 | 92.235 | 0.029 |
| *Spaceflight with gravity control vs. ground control in shoot tissue* | -613.833 | 126.545 | 1.23e-6 |
| *Spaceflight with gravity control vs. ground control in root tissue* | 210.625 | 134.222 | 0.117 |
| *Spaceflight vs. ground control in shoot tissue* | 641.333 | 107.284 | 2.26e-9 |
| *Spaceflight vs. ground control in root tissue* | -190.625 | 122.527 | 0.120 |

**Supplementary Table S3. Comparing the treatment groups, full list of estimates.** Interaction effects were found between spaceflight stress conditions and tissue type. All significant interacting variables (gravity level, tissue type, and non-microgravity spaceflight stress (namely space radiation)) must be incorporated in the comparisons to draw the appropriate conclusion. The non-microgravity spaceflight stress (namely space radiation) impacted the number of variants called in gravity and shoot tissue. The estimates highlighted in green are the significant results that incorporate all interactions appropriately.

PREPRINT
